# Unsupervised manifold learning of collective behavior

**DOI:** 10.1101/2020.03.23.003244

**Authors:** Mathew Titus, George Hagstrom, James R. Watson

## Abstract

Collective behavior is an emergent property of numerous complex systems, from financial markets to cancer cells to predator-prey ecological systems. Characterizing modes of collective behavior is often done through human observation, training generative models, or other supervised learning techniques. Each of these cases requires knowledge of and a method for characterizing the macro-state(s) of the system. This presents a challenge for studying novel systems where there may be little prior knowledge. Here, we present a new unsupervised method of detecting emergent behavior in complex systems, and discerning between distinct collective behaviors. We require only metrics, *d*^(1)^, *d*^(2)^, defined on the set of agents, *X*, which measure agents’ nearness in variables of interest. We apply the method of diffusion maps to the systems (*X, d*^(*i*)^) to recover efficient embeddings of their interaction networks. Comparing these geometries, we formulate a measure of similarity between two networks, called the map alignment statistic (MAS). A large MAS is evidence that the two networks are codetermined in some fashion, indicating an emergent relationship between the metrics *d*^(1)^ and *d*^(2)^. Additionally, the form of the macro-scale organization is encoded in the covariances among the two sets of diffusion map components. Using these covariances we discern between different modes of collective behavior in a data-driven, unsupervised manner. This method is demonstrated on empirical fish schooling data. We show that our state classification subdivides the known behaviors of the school in a meaningful manner, leading to a finer description of the system’s behavior.

**Author summary:** Many complex systems in society and nature exhibit collective behavior where individuals’ local interactions lead to system-wide organization. One challenge we face today is to identify and characterize these emergent behaviors, and here we have developed a new method for analyzing data from individuals, to detect when a given complex system is exhibiting system-wide organization. Importantly, our approach requires no prior knowledge of the fashion in which the collective behavior arises, or the macro-scale variables in which it manifests. We apply the new method to an agent-based model and empirical observations of fish schooling. While we have demonstrated the utility of our approach to biological systems, it can be applied widely to financial, medical, and technological systems for example.

## Introduction

Collective behavior is an emergent property of many complex systems in society and nature [1]. These behaviors range from the coordinated behaviors of fish schools, bird flocks, and animal herds [2–4], to human social dynamics such as those of traders in stock markets that have led to bubbles and crashes [5] in the past, and opinions on social networks [6]. Collective behavior amplifies the transfer of information between individuals and enables groups to solve problems which would be impossible for any single group member alone [7]. Furthermore, manifestations of collective behavior in both social and ecological systems regularly influence our individual welfare [8–10] through political and economic instability, disease spread, or changes in social norms. Building a better classification and understanding of collective behavior and collective states is crucial not only to basic science, but to designing a stable future.

A key challenge in the study of complex systems is to identify when collective behavior emerges, and in what way it manifests [11, 12]. Statistical physicists first studied emergence in complex systems in the 1800s where they began to derive the physical laws governing the macroscopic behavior of systems with an extremely large number of degrees of freedom from the interactions between the microscopic, atomistic components of the system. Remarkably, there usually exist a small number of macroscopic variables or order parameters that accurately describe macroscale dynamics for numerous systems, and a small number of relationships between these variables characterize the different states or behavioral regimes of the system. In many systems, the key macroscopic variables are well known from either first principles or empirical study. However, there are many social, ecological, and even physical systems which elude simple description by macroscopic variables. Furthermore, we continually encounter novel systems for which we have no prior knowledge about multiscale dynamics, or models that give rise to unknown group dynamics [13]. In these cases, it is a non-trivial task to identify useful variables (at particular scales of organization) which define the possible emergent behaviors.

Machine learning and other statistical methods have emerged as key tools for finding macroscopic descriptions of complex systems with many degrees of freedom, in the physical, ecological, and social sciences. The application of these tools has been driven by high resolution observations of microscopic degrees of freedom (particularly in ecological and social systems), increases in computing power, and the development of new algorithmic tools. Dimensionality reduction, clustering algorithms, and other unsupervised learning algorithms have been particularly successful in discovering macroscopic variables and behavioral regimes from microscopic data on complex systems [14]. The paramagnetic/ferromagnetic phase transition in the classical, 2D Ising model has been discovered by algorithms ranging in complexity from principal component analysis to variational autoencoders [14–16]. However, more complex types of collective phenomenon, such as topological phases, have not been successfully identified using algorithms like PCA, which assume that the data has a linear structure. In these more difficult cases, algorithms which can detect nonlinear structures such as diffusion maps have shown much greater promise. For example, diffusion maps have been used to find the Kosterlitz-Thouless phase transition in the XY-model from condensed matter physics [17]. The success of machine learning algorithms for physical problems suggests that machine learning could be of great practical use for the study of collective phenomena in social and ecological systems, which are usually much less amenable to theoretical analysis, and for which there often exists a significant quantity of data at the individual level.

Here we have developed an approach to analyzing collective behavior in biological systems based on dimensional reduction tools from manifold learning [18], specifically diffusion maps [19]. Using diffusion maps to formulate multiple data-driven coordinate systems, we compare their large-scale structures to find latent global relationships between different variables observed at the level of individual agents. We call this the map alignment between the two coordinate systems. When two variables serve to globally organize the data along similar lines, their map alignment is high. One can easily calculate the expected alignment between unrelated and independent variables, so this gives us a test that detects a system’s emergent behaviors but requires no prior knowledge of them. Moreover, the ways in which the coordinates are correlated leave a signature for the relationship between the macro-scale structures, and one may use this to distinguish various system-wide behaviors. We apply this tool to modeled data produced from simulations of birds flocking, and to empirical data on the activity of fish in a school. Using the micro-scale data to cast the agents’ network of interaction into multiple geometries, each evolving over time, we both detect the emergence of macro-scale organization as well as perform a meaningful classification of distinct modes of organization, identifying the macro-scale states of the system. Importantly, this analysis is based purely on changes occurring at micro-scales, and makes no assumptions about which macro-scale variables are important. This means that for any novel system of study, where there is a paucity of knowledge regarding which macro-scale variable to measure and track, one could use this approach to classify and observe the dynamics of the system at different levels of organization.

## Materials and methods

### Diffusion Maps

We begin with the basic definitions required to construct a diffusion map. This requires a set of data points, *X*, and a metric on those points, *d*. The data may be composed of many observables, making each *x*_*k*_ ∈ *X* a vector in a space of potentially very high dimension. Essentially we employ the algorithm in [20], with *α* = 1 and *ϵ* = 1. This involves

i. constructing a similarity measure, *K*(*x*_*i*_, *x*_*j*_), between pairs of data points,
ii. normalizing *K* to create a Laplacian operator *Â* on the data,
iii. computing eigenvector-eigenvalue pairs {(*ϕ*_*k*_, *λ*_*k*_)} for the Laplacian, and
iv. constructing the diffusion map 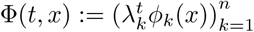.

The choice of similarity measure in (*i*) is highly system-dependent, for agent-wise interactions may manifest themselves in many different ways and so require different metrics.

In (*ii*), the choice of normalization in constructing *Â* is subtler, and based on a choice of Laplace operator; this is not sensitive to the data under observation, but rather reflects an assumption of Manifold Learning, that the data points of *X* are drawn from a manifold embedded in euclidean space. For rigorous results on when such a manifold exists for a given dataset, see [21].

Given *n* points *X* = {*x*_1_, …, *x*_*n*_}, we construct an *n* × *n* matrix of pairwise distances, *D* = (*D*_*i,j*_) := (*d*(*x*_*i*_, *x*_*j*_)), associated to a metric *d*. Writing *k*(*x*) for the *k*^th^ nearest neighbor (kNN) of *x* ∈ *X*, let *K*(*x, y*) be the gaussian kNN (GkNN) kernel defined by

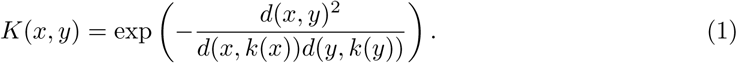

This kernel could be replaced with any other, as another function may better reflect the user’s impression of the similarity between two agents a distance *d*(*x, y*) apart. This particular choice is studied in [22], and serves to allow pairwise distances to scale according to their local geometries. One could consider this the standard gaussian kernel applied to the modified distance function 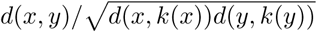. The more sparse the point cloud (*X, d*) is near *x* or *y*, the smaller this distance becomes. This kernel also has the desirable property that it is scale-free, in that multiplying all distances by a fixed constant does not change the kernel *K*.

Let *σ* denote a row sum operator: if *A* is an *m* × *n* matrix, let *σ*(*A*) be the *m*-vector whose *i*^th^ component is 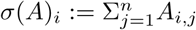 We next calculate an affinity matrix *A* = (*A*_*i,j*_) := (*K*(*x*_*i*_, *x*_*j*_)), its row sums *σ*(*A*), and construct the matrix operator *A*′ via

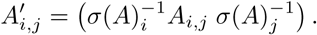

Normalizing by the row sums in this fashion is a particular choice of Laplace operator on the graph whose vertices represent the *n* agents and whose weighted edges are given by the entries of 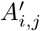 (see panel A of Figure 2). This operator converges pointwise to the Laplace-Beltrami operator in the case that the points *X* are drawn from a probability distribution on a manifold (see [20]). We then define the Markov operator *Â* = (*Â*_*i,j*_) by setting the row sums of *A*′ to one:

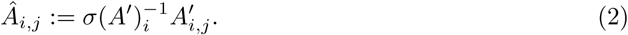

**Fig 1.**
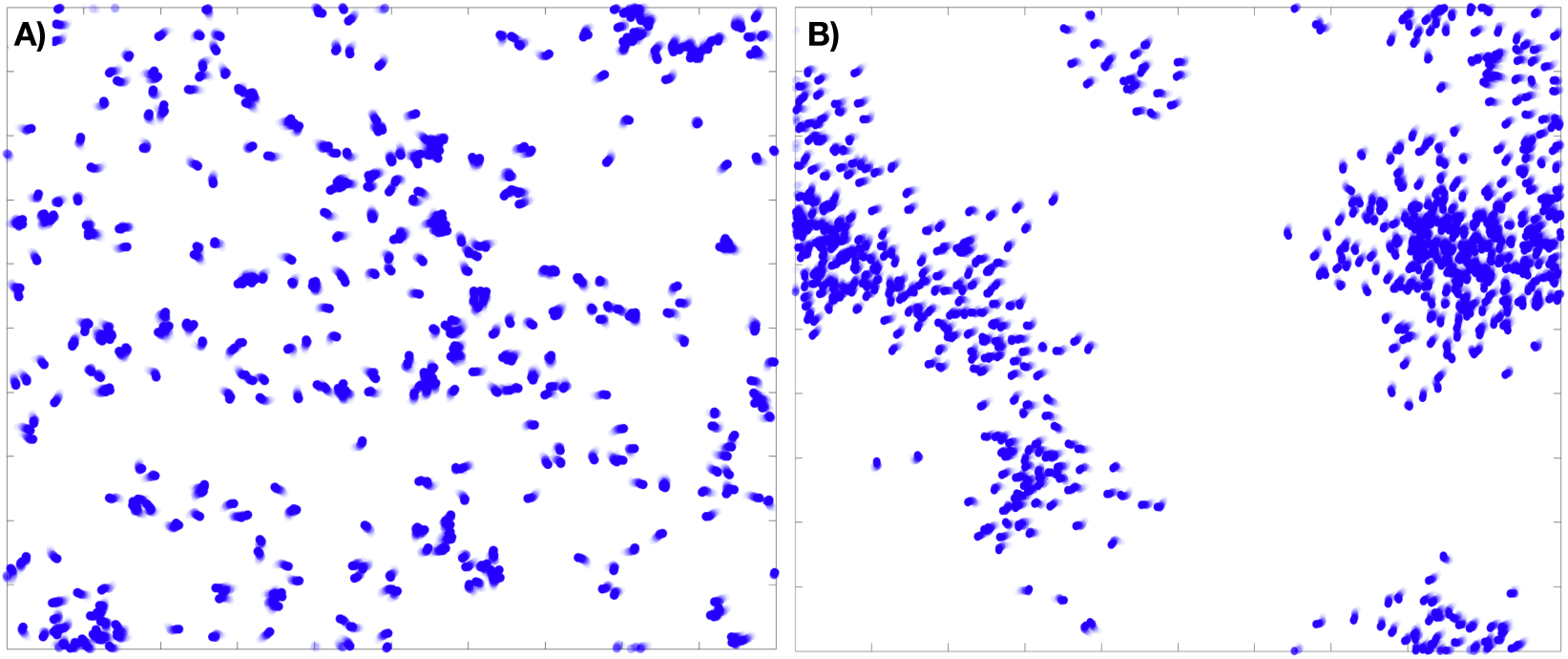
Examples of an 800 agent flocking model on the torus, with and without long-range order, that is, in both incoherent (A) and coherent (B) macro-states.

**Fig 2.**
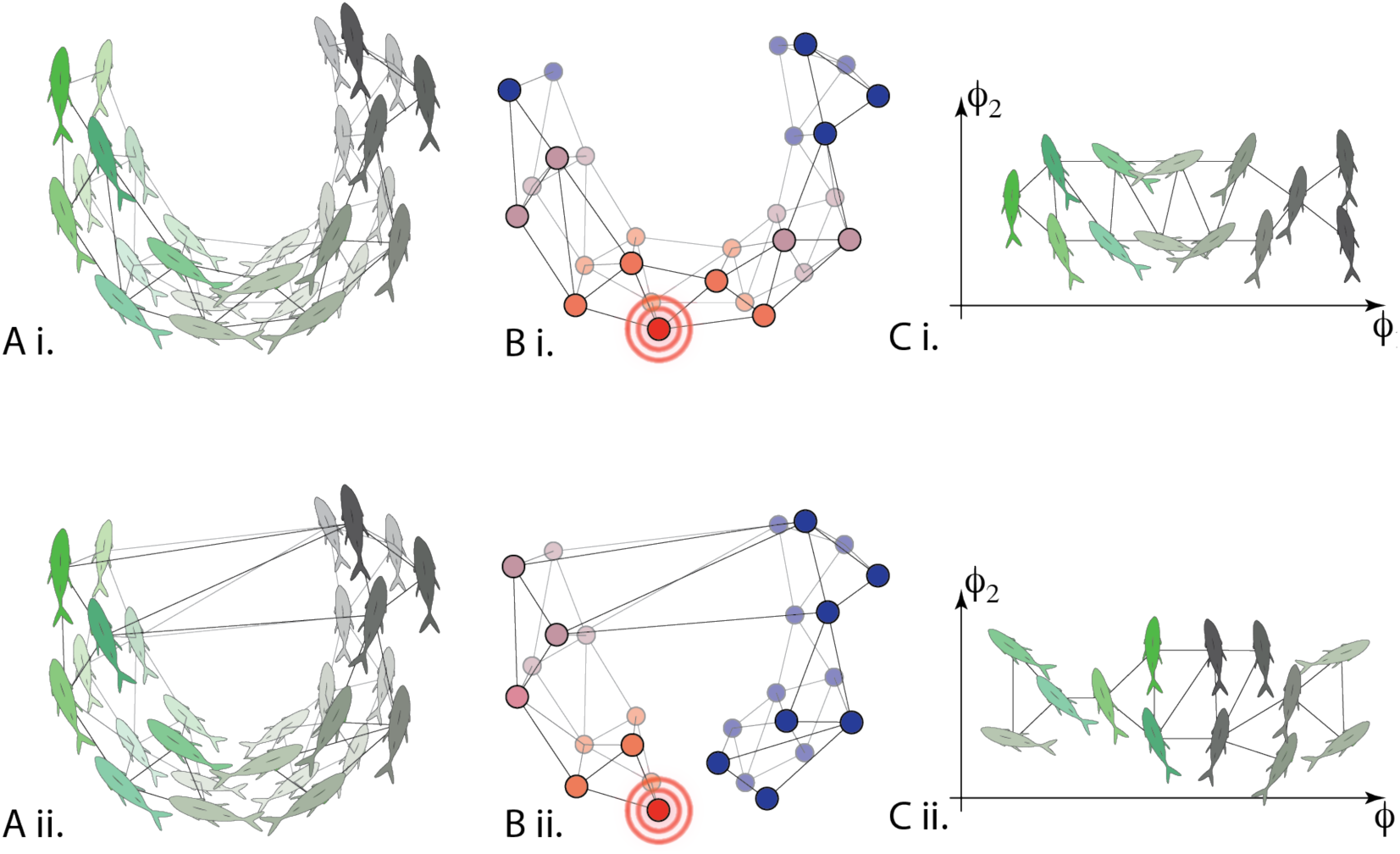
Conceptual illustration of diffusion maps applied to data on fish schooling. A) For the same time period, different affinity operators can be calculated. These affinities could be correlations in simply spatial proximity (Ai) or correlations in velocities (Aii) for example. The operator is used to describe a network of agents. B) Once calculated, the heat diffusion operator allows us to quantify the geometry of the network and with this in hand C) diffusion map coordinates may be used to efficiently embed the fish in a low-dimensional space. Doing this for different metrics / affinities allows us to quantify the alignment between various diffusion map coordinate systems, which reveals changes in macro-scale behavior.

This is the operator of primary interest to us, representing the evolution of a random walk on the graph, where the probability of transitioning from point *x*_*i*_ to *x*_*j*_ is given by *Â*_*i,j*_. This is visualized in panel B of Figure 2 as heat propagating through the network. From the spectral structure of *Â*, written 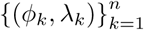 with *ϕ*_*k*_ : *X* → ℝ the unit eigenvectors and *λ*_*k*_ ∈ ℝ the eigenvalues, we construct the diffusion map,

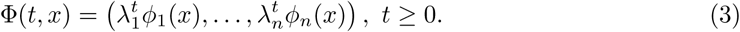

The usefulness of this decomposition of the data (*X, d*) is that it gives an embedding of the data which reflects the *diffusion distance, D*_*t*_, defined as the *L*^2^ difference between the random walk’s distributions when initiated from site *x* and site *y* after *t* time has passed:

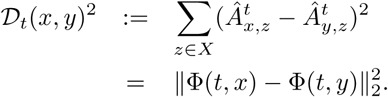

In essence, *𝒟*_*t*_ captures the geometric structure of the data by measuring the (dis)similarities of trajectories of random walks evolving on the data points from different starting points. The diffusion map is an isometry of (*X, D*_*t*_) into ℝ^*n*^, with the remarkable benefit that truncating Φ(*t, ·*) to its first *k* coordinates gives the optimal *k*-dimensional embedding of *X* into ℝ^*k*^ (envisioned in panel C of Figure 2). These truncated maps 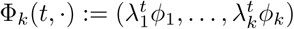 are utilized below for detecting macro-scale relationships among variables.

We remark that though increasing *t* decreases the resolution of small scale features in the embedding, in practice we take *t* = 1 throughout and define *ψ*_*k*_(*x*) = *λ*_*k*_*ϕ*_*k*_, so that

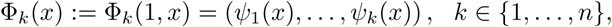

and 𝒟 := 𝒟_1_.

### Map Alignment

In this section we formulate a statistic to measure the degree to which the macro-scale organization of two observables measured on the micro-scale level are related. Let *X* = {*x*_1_, …, *x*_*n*_} be the collection of data for the *n* interacting agents making up the system, and let *f*_1_ and *f*_2_ be two vector-valued observables, such as position or velocity, defined on each agent,

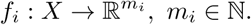

Suppose we are given metrics *d*_*i*_ : *X* × *X* → ℝ between agents depending only on the observed variable *f*_*i*_, such as

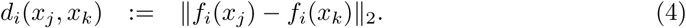

In practice these are defined by the user, and a cogent choice will depend upon their knowledge of the agent-level interactions of the system. Using the methods of the previous subsection, we devise a pair of Markov operators, *Â*^(*i*)^ (taking *i* = 1, 2), their associated eigenstructures, 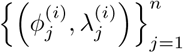, and the corresponding diffusion maps, 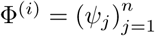.

For each *i* = 1, 2 the diffusion coordinates 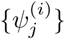are ordered by decreasing contribution to the global organization of the network *X* under the heat operator *Â*^(*i*)^. Therefore if there is a coherent large-scale organization on *X* manifest in the observables *f*_*i*_, then the leading diffusion coordinates of Φ^(1)^ should be determined in some manner by the leading coordinates of Φ^(2)^. To confirm this, we expand the unit vectors 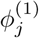 in the subspace generated by the top *k* diffusion coordinates of *Â*^(2)^, i.e. we project the eigenvectors into the subspace spanned by 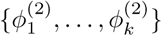 via the linear operator

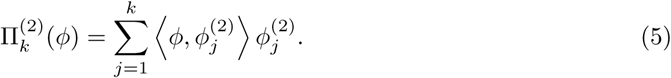

Then 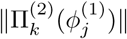 is a measure of how well the coordinate 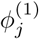 associated to *f*_1_ is correlated with the large-scale network structure induced on *X* by the variable *f*_2_. One attempts to choose *k* in a manner that ensures that the subspace spanned by 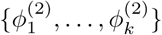 encompasses the relevant macro-scale coordinates induced by *f*_2_ on *X*, but without *k* becoming so large that we begin to recover a significant portion of *ϕ* simply due to the large dimension of the image space. For example, if we take *k* = *n* the projection 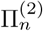 is the identity, simply expressing *ϕ* in the basis 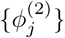. However, taking *k* ≪ *n* we recover only the portion of the vector which can be expressed as a linear combination of the top *k* diffusion coordinates of Φ^(2)^. We return to this issue below.

Now we derive the expected projection size under a null hypothesis that the two variables *f*_1_ and *f*_2_ are entirely unrelated. Given the pair of diffusion bases, we fix 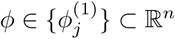 and calculate the distribution of the random variable 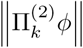 assuming that the *k*-dimensional range, Span 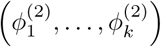, is chosen uniformly at random from the set of all *k*-dimensional subspaces. One can show, and with a little thought it is clear, that this is equivalent to asking for the magnitude of a random unit vector of ℝ^*n*^ projected onto its leading *k* coordinates. When the vector is sampled uniformly at random from {*ψ* ∈ ℝ^*n*^ : ‖*ψ*‖_2_ = 1}, the quantity 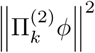 is a beta distributed random variable with parameters (*n* − *k*)*/*2 and *k/*2. Recognizing this, one can calculate the expectation and variance to be *k/n* and 2(*n* − *k*)*k/*(*n*^3^ + 2*n*^2^), respectively. Normalizing the projection size accordingly, we arrive at the statistic

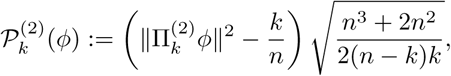

which quantifies how well we can express the information of *ϕ* using only the random subspace generated by 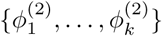.

Again, the leading diffusion coordinates of Φ^(*i*)^ are data-driven macro-scale variables, expressing the large-scale organization of the network as viewed through *Â*^(*i*)^. If the data obeys a (possibly nonlinear and multi-valued) relationship between *f*_1_ and *f*_2_ on the macro-scale, then it should be evidenced by a relationship between the bases 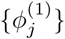 and 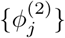. Note that the observables *f*_*i*_ are used only in providing a metric on *X*. One can perform an identical analysis on the data given only the metrics *d*_1_ and *d*_2_. In the context of this paper we will assume, however, that the metrics come from some micro-scale variable defined on *X*, as this is the approach most useful in analyzing a new complex system.

Let 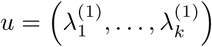 be the vector of decreasing eigenvalues of *Â*_1_, and take 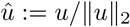. Then the entries of *û* give a normalized notion of the energy of each 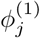, and so gives a measure of how important each mode 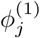 is to the macrostructure derived from *f*_1_. Finally, we define the **map alignment statistic** to be the sum of the 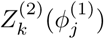, weighted by the contribution of 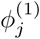 to the large-scale organization of *f*_1_:

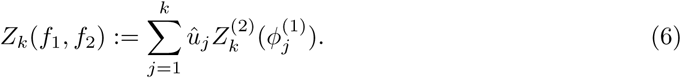

We remark that this statistic is not in general symmetric; *Z*_*k*_(*f*_1_, *f*_2_) ≠ *Z*_*k*_(*f*_2_, *f*_1_). The MAS *Z*_*k*_(*f*_1_, *f*_2_) is a measure of how much of the macrostructure of *f*_1_ is reflected in the *f*_2_-based diffusion coordinates. For example, the spectrum of the *Â*^(*i*)^ may indicate that the top three diffusion coordinates of 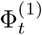 carry a greater proportion of the macro-scale information of *Â*^(1)^ than the top three coordinates of 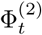 do for *Â*^(2)^; in this case one would expect that *Z*_3_(*f*_1_, *f*_2_) > *Z*_3_(*f*_2_, *f*_1_).

### Informative choice of *k*

Here we briefly discuss how to choose *k* so that *Z*_*k*_ gives an accurate measure of the degree to which Φ^(1)^ is determined by 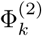. Recall that the head of the spectrum of *Â* is sensitive to large-scale changes, but the bulk of the spectrum is associated to high frequency eigenfunctions, capturing small-scale organization often attributable to noise or local idiosyncrasies in the data. As a result, given training data or simulated data one may simply estimate a value of *k* after which the variance explained by the *j*^th^ eigenspace, 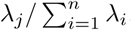, is approximately independent of the macrostate of the system. That is, choose *k* so that for *j* > *k*, the value *λ*_*j*_ makes up a fixed fraction of the sum of the eigenvalues.

Alternatively, in using empirical data we found the bulk of the spectrum often follows an approximate power-law decay. As a result, one may take *k* to be the smallest integer such that for *j* > *k, λ*_*j*_ ≈ *Cj*^−*β*^ for positive constants *C, β*.

These are necessarily rough guidelines, and in practice one often finds a range of plausible choices for *k*. However, when increasing *k* the additional summands of 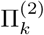 must eventually become essentially independent 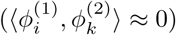 of the leading elements of 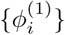. It follows that after a point, a larger value of *k* has little influence on the computed MAS, *Z*_*k*_(*f*_1_, *f*_2_). Therefore an overestimate of *k* has very little effect on the results, so it is advised that in practice one err on the side of larger choices of *k*.

### Unsupervised Macrostate Classification

If the system under study exhibits multiple emergent behaviors relating the two variables *f*_1_ and *f*_2_, then coherence alone is not enough to identify system behavior. In passing to the norm of the vector 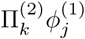, we clearly give up a significant amount of information. In order to distinguish multiple coherent states, we inspect the individual dot products 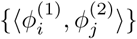 that comprise the map alignment statistic.

However, to remove the effects of the trailing diffusion coordinates, which correspond to smaller scale variations in the geometry of (*X*, 𝒟^(*i*)^), we consider the sequence of inner products given by

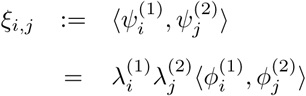

with 1 ≤ *i, j*, ≤ *k* ≪ *n*. That is, we collect the covariances between coordinates of the truncated diffusion maps 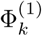 and 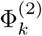. Given time series of micro-scale observations, this allows us to characterize each frame by the sequence (*ξ*_*i,j*_)1≤*i,j*≤*k*. We will add an argument to indicate the timestep *t*, as *ξ*_*i,j*_(*t*), when necessary.

While the diffusion coordinates provide a natural organization of the data, the individual components of the map are eigenvectors, and so are computed via an iterated maximization problem: given the linear operator *A*, the *i*^th^ eigenvector is given by

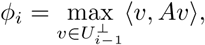

where 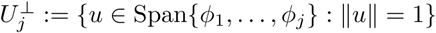. Such maximization problems are prone to discontinuities, in the sense that there exist, for any threshold *ε* > 0, linear operators *A* and *B* on ℝ^*n*^ with *i*^th^ eigenvectors *ϕ*_*i*_ and *η*_*i*_, respectively, such that ‖*A* − *B*‖ < *ε*, while ‖*ϕ*_*i*_ − *η*_*i*_‖> 1*/*2 for all 1 ≤ *i* ≤ *n*. This means that the individual indices of the *ξ*_*i,j*_ may not correspond to a persistent structure in the data.

Therefore, for each collection {*ξ*_*i,j*_ : 1 ≤ *i, j* ≤ *k*} we define the map *σ* : [*k*] × [*k*] → [*k*^2^] so that the components of (|*ξ*_*σ*(*i,j*)_ |) are ordered by decreasing magnitude. (Here [*k*] denotes the first *k* positive integers.) This removes the ambiguity in both index and sign for these products. We call the resulting vector the *f*_1_*f*_2_-**covariance vector** (at time *t*), abbreviated as *ξ* (resp. *ξ*(*t*)). One may then use these vectors as a fingerprint for the macro-scale relationship exhibited between *f*_1_ and *f*_2_ at a given time.

Figure 3, panel A, gives an example of a toy flocking system comprised of 50 agents, exhibiting a circling behavior at time *t*_1_ and a (mostly) aligned state at time *t*_2_; its top four diffusion coordinates associated to both Φ^(1)^ and Φ^(2)^ are displayed for each time. (The choice of affinities *Â*^(1)^ and *Â*^(2)^ here are the same as in the flocking model below, being based on spatial proximity and velocity, respectively.) In panel B we see the absolute values of the inner products of the diffusion coordinates, where the matrix (represented by a color plot) has its (*i, j*)-entry given by 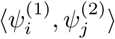. Finally, in panel C we see the covariance vectors at times *t*_1_ and *t*_2_.

**Fig 3.**
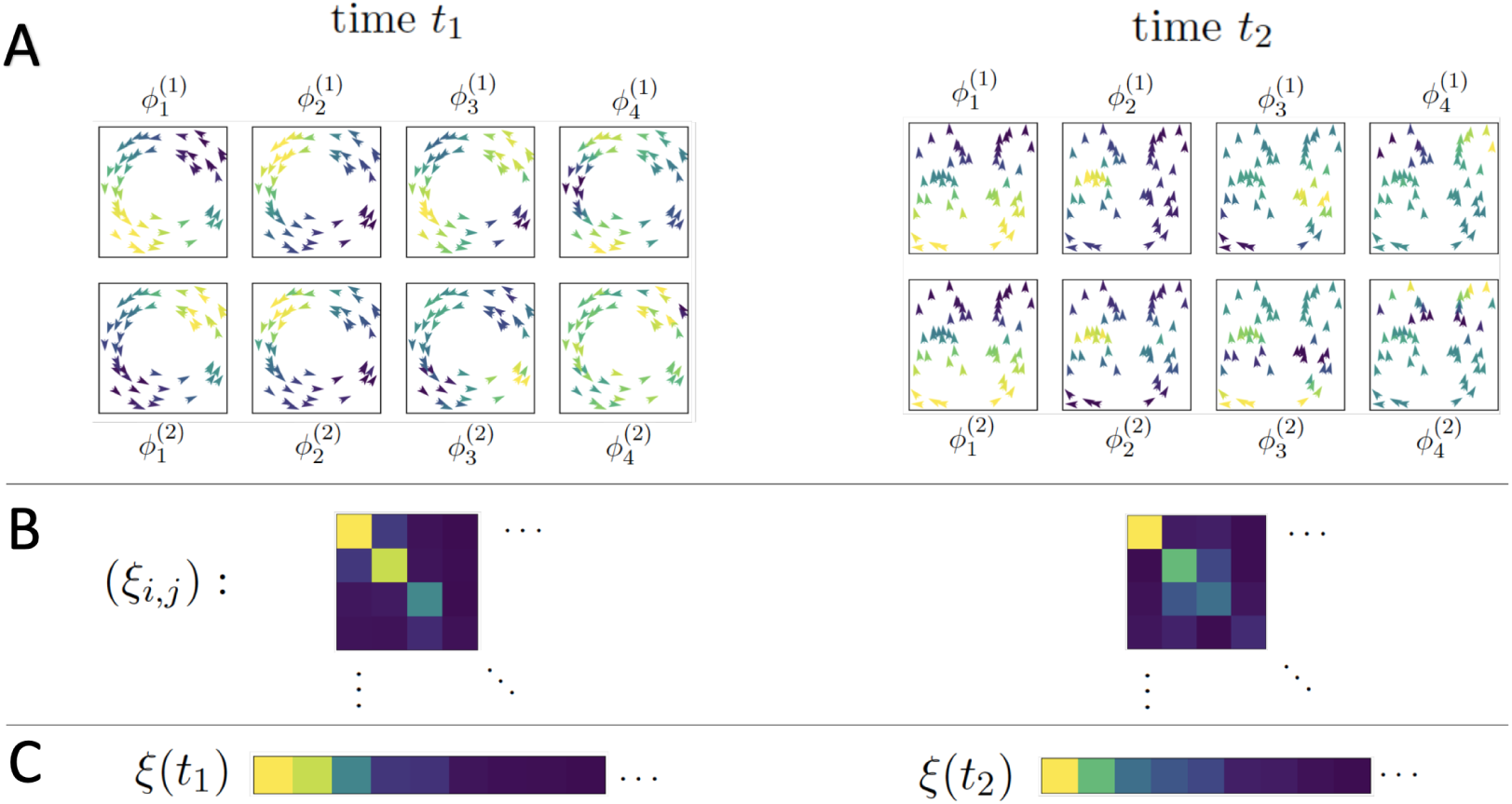
A toy system of 50 agents; the left and right sides of the figure represent observations of the system at two different times when the system is in two different organized states. (A) Agents in each subpanel are colored by the indicated diffusion coordinate, (B) A color plot of the matrix (*ξ*_*i,j*_) with yellow indicating larger values, blue indicating smaller values, (C) The values of the above inner product matrix, ordered by absolute magnitude; these are the vectors used to classify the system’s regimes.

Covariance vectors allow us to embed the activity of the agents in a shared space, 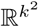, and perform clustering analyses to group similar macro-scale relationships between *f*_1_ and *f*_2_ over time. In the next section we will perform a boilerplate clustering analysis to distinguish between distinct coherent group structures found in a data set of golden shiner schooling behavior, a task for which the map alignment statistic is insufficient on its own.

## Results

### Analysis of Bird Flocking Simulations

As a first test of our method we explored a well studied agent-based model of flocking behavior [23]. The model consists of a set of *N* agents, {*x*_*j*_}, moving on the torus *T* ^1^ = *S*^1^ × *S*^1^, each with position *p*_*i*_ and velocity determined by the angle *θ*_*i*_ and the magnitude *v*_*i*_. In this model each agent moves with a fixed speed, *v*_*i*_ = *v*_*j*_ for all *i* and *j*, so we simply write *v* for this speed.

Each agent’s direction is updated at each time step by the average local direction plus a noise term, a normal random variable distributed as *N* (0, *ϵ*^2^). Here local refers to the agents within a certain distance *r* of the focal agent. With all other parameters of the model fixed, the coupling parameter *r* governs the degree to which the system self-organizes. In the hydrodynamic limit, long-range order emerges for *r* above some critical value *r*_*c*_(*v, ϵ*) > 0, meaning that the mean velocity of the flock is nonzero: 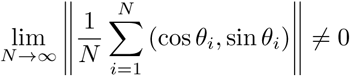. The direction of group travel is chosen randomly, and breaks the symmetry of the system. See the Supplementary Information (SI) for details of the model, video, and pseudocode.

In the discrete time, finite-agent setting we observe the mean velocity of the flock, ⟨*v*_*j*_⟩ = (*v* cos⟨*θ*_*j*_⟩, *v* sin⟨*θ*_*j*_⟩), taking this as our ‘ground truth’ measurement of the level of emergent, coherent behavior in the system. As *r* increases past *r*_*c*_(*v, ϵ*), the mean velocity (in the large *N* limit) transitions from zero to a nonzero value. However, this shift occurs continuously; as *r* grows we find the steady-state mean velocity of the flock increases towards its maximum value. Even with *r* = 0 the global mean velocity will have magnitude 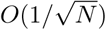 and suffer fluctuations, so determining whether a preferred flock-wide direction of motion has been chosen in the finite case makes little sense. Instead, below we compare the value of *Z*_*k*_(*p, v*) to ⟨*v*_*j*_⟩ to determine how well the MAS detects large-scale organization.

We chose two metrics to define a pair of time-evolving graphs as follows: Since we are interested in flocking behavior, we let *d*^(1)^(*x*_*i*_, *x*_*j*_; *t*) be the euclidean distance between the *i*^th^ and *j*^th^ agents at time *t*, while *d*^(2)^(*x*_*i*_, *x*_*j*_; *t*) denotes the kernel-smoothed *L*^2^ norm of the two agents’ relative speed profiles. (See (*S*3), (*S*4) and (*S*5) for the explicit definitions of the metrics used.) In particular, *d*^(1)^ captures the spatial information in the flock, while *d*^(2)^ measures the degree that agents are moving in concert over a short (on the order of 10 time steps) recent period.

We then applied diffusion maps to the resulting sequence of graphs and calculated the map alignment between the two variables, *p* and *v*. The choice of subspace size to use, *k* = 10, was empirical. We observe that at various levels of organization the variance explained by the first five modes (i.e. the sum of the magnitudes of the first four components of *û*) is highly variable, while the variance explained by the higher modes 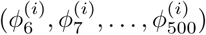 shows little dependence on system organization, though we decided to increase *k* to 10 to be conservative. This suggests that the macro-scale information is well captured by 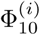, while components 11 through 500 are largely accounting for noise and small-scale features of *Â*^(*i*)^. Note, however, that one may take *k* larger than 50 without any qualitative change in the analysis.

In Figure 4 below we plot the map alignment statistic *Z*_10_(*p, v*) (blue) calculated from two sets of 100 independent simulations of the flocking model with 500 agents and fixed parameters (*ϵ* = *π/*5, *v* = 1*/*320, see SI). In the left panels the coupling parameter *r* is held at a low value (*r* = *r*_0_ = 0.0065) for 800 time steps before increasing linearly to a high value over 400 steps and remaining at that value (*r* = *r*_1_ = 0.06) for the rest of the trial; see the lower panel of Figure 4(A). The mean velocity ⟨*v*_*j*_⟩ of the flock, averaged over the 100 trials, is plotted in orange in the upper panel.

**Fig 4.**
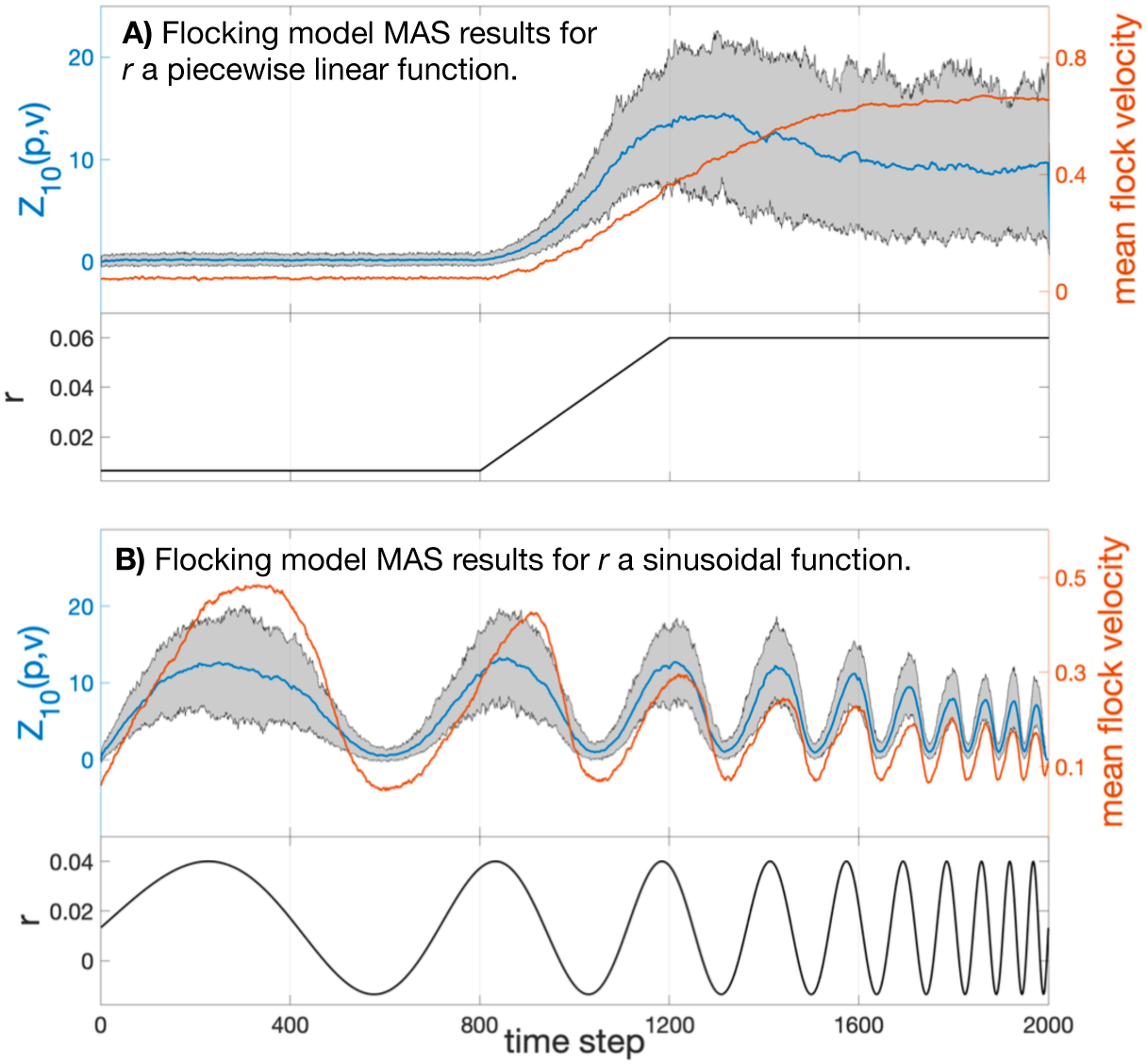
Top: Map alignment statistic *Z*_10_(*p, v*) averaged over 100 flocking simulations of 2000 time steps (blue). The grey region denotes the 90% empirical confidence interval of *Z*_10_(*p, v*). The mean velocity ⟨*v*_*j*_⟩ of the flock is averaged over the 100 trials and plotted in orange. Bottom: The model’s coupling constant *r* plotted over time. Notice that in (B) the MAS does not suffer the same hysteresis that the flock’s speed does.

One can see that the statistic *Z*_10_(*p, v*), measuring the dependence of position diffusion coordinates on the velocity diffusion coordinate system, has a strong correspondence with the velocity correlation statistic. That is, large-scale organization was clearly detected and quantified using only the two micro-scale inter-agent distance time series.

In a second trial we explored how quickly map alignment and long-range correlations vary in response to change in the micro-scale dynamics of the agents. This is done by allowing *r* to vary sinusoidally between *r*_0_ and *r*_1_, with increasing frequency. The black line plotted in the lower panel of Figure 4(B) shows the coupling parameter as a function of time; in the upper panel the orange line shows the magnitude of the mean velocity of the flock, and the blue plot gives the mean value of *Z*_10_ across trials, with the filled area indicating the 90% confidence interval for *Z*_10_.

Clearly *Z*_10_(*p, v*) indicates strongly the presence or absence of correlation. The map alignment statistic shows both a faster response to changing *r* and suffers from less attenuation as the speed of parameter change increases. Times *t* = 800 to 1000 especially highlight how the drop in *Z*_10_ is a precursor to the value of the macrovariable ⟨*v*_*j*_⟩ falling.

In this case and the previous case, we find we are able to track the transition from unorganized motion to flocking by the alignment between the velocity diffusion coordinates, 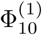, and the position diffusion coordinates, 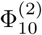.

### Analysis of Empirical Fish Schooling Data

As an empirical application we use position and velocity data for a school of golden shiners (*notemigonus crysoleucas*); details of how the data were collected can be found in the Supplementary Information. These fish were studied in [24, 25]. In [25] the authors found that the school, constrained to an essentially two dimensional environment, would generally be found in one of three macro-states defined according to the school’s collective polarization, *O*_*p*_, and rotation, *O*_*r*_: milling (circling), swarming (disordered, stationary), or polarized (translational). See Figure S1 for example frames of these behaviors. Writing *v*(*x*_*i*_) for the velocity of fish *i* and *p*(*x*_*i*_) for its position, the chosen macrovariables are defined as

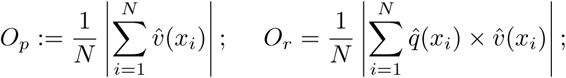

where *q*(*x*_*i*_) := (*p*(*x*_*i*_) − *c*) is the vector from the school’s centroid, 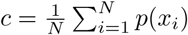, to the position of fish *i*. Hats indicate unit vectors and × indicates the cross product. One can then demarcate the behaviors as follows:

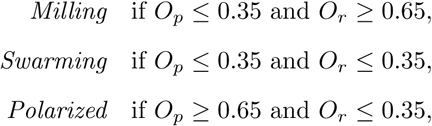

with all other (*O*_*p*_, *O*_*r*_) values considered to be *transitional* states. In the lower panel of Figure 5 the variable *O*_*p*_ is plotted for the 5000 frames used in this study; the coloration of the plot indicates the school’s state as defined here (red for milling, yellow for swarming, dark green for polarized, and grey for transitional). The remainder of this section demonstrates that the unsupervised tools developed above are capable of discerning data-driven macrostates corresponding to, and even refining, those defined in [25].

**Fig 5.**
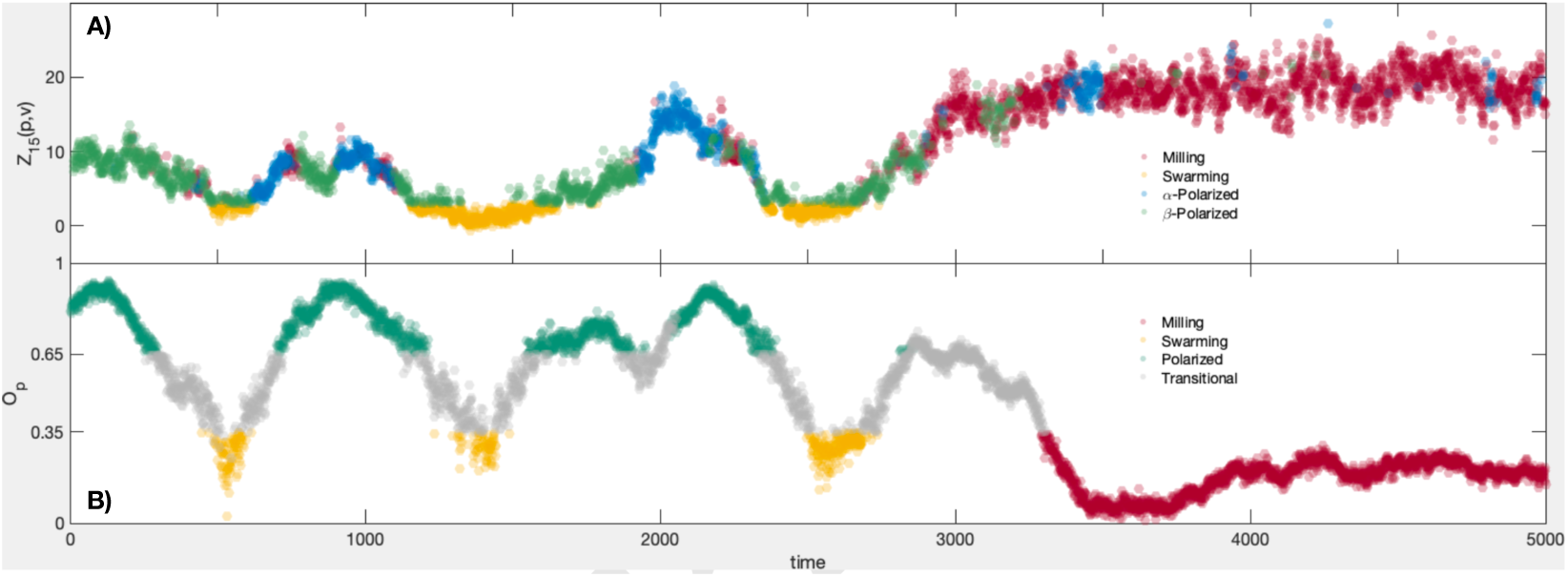
Statistics calculated from 5000 frames of the golden shiner dataset. A): Map alignment *Z*_15_(*p, v*) calculated over 5000 video frames. The plots are colored according to our classification of macro-scale state; milling = red, swarming = yellow, *α*-polarized = blue, *β*-polarized = green. B): The group polarization macro-variable *O*_*p*_ calculated from the same 5000 frames. The coloring shows the states according to [24]; milling = red, swarming = yellow, polarized = teal, unclassified/transitional = grey.

For the study of fish schooling, as with the simulated data above, we built a pair of distance matrices at each time step. The first was calculated from the euclidean distances between agents (individual fish), *d*^(1)^, defined as before. The second, *d*^(2)^, using the velocity data only. In this case, our goal was to capture any interaction between agents, either instantaneous or at a lag of ≥ 1 time step due to one fish leading another. Therefore we define *d*^(2)^(*x*_*i*_, *x*_*j*_; *t*) as in the previous section (a weighted *L*^2^ norm of the differences in the agents’ recent velocity profile), but now allow one of the two agent’s velocity profiles to begin at an earlier time step, *t* − *ℓ* with 0 ≤ *ℓ* ≤ 60. We take *d*^(2)^(*x*_*i*_, *x*_*j*_; *t*) to be the minimum of this collection of distances, as this choice of time translation gives the best alignment of the two fishes’ trajectories.

As with the flocking model, the first diffusion map’s leading coordinates identify the large groups of fish who form spatially proximate clusters; the leading diffusion coordinates associated to *d*^(2)^ highlight groups whose velocity profiles are (up to a 2s, or 60 frame, translation in time) quite similar. The map alignment between these two systems, *Z*_15_(*p, v*), is plotted in the upper panel of Figure 5. When the fish are swarming (see Figure S1, center column) we see that these two coordinate systems are unrelated, and that *Z*_15_(*p, v*) is correspondingly low.

Here again, the choice of *k* = 15 is driven by observing that outside of the top fifteen entries of *û* = *u*/‖*u*‖ for 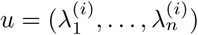, *i* ∈ {1,2}, the spectral decay appears to follow a power law for most frames (though this observation does break down for some frames when the eigenvalues become small enough). The results below prove to be robust to varying *k* from 10 to 25. Therefore the span of 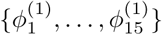 contains the majority of the structure within the data, while *k* = 15 is not so large that the signal in *Z*_*k*_ becomes diluted. The two coherent behaviors we would like to separate are the polarized, or linear, motion (right column in Figure S1) and milling, or circling, motion (left column in Figure S1). Both states exhibit high levels of organization, and we must use a sharper tool to distinguish them.

To do this we employ the covariance vectors *ξ*_*i,j*_, as defined in the Unsupervised Macrostate Classification section above, over the course of 5000 frames of video (see the Supplementary Information (SI) for more information on the dataset). First, those states with *Z*_15_(*x, v*) < 3 are considered to exhibit incoherent behavior and to have no macro-scale dynamic, so we classify them as swarming frames and remove them from the set of frames to be classified. Then for each time *t* we compute the covariance vector *ξ*(*t*) = (|*ξ*_*σ*(*i,j*)_(*t*)|). We cluster the resulting vectors using the *k*-means algorithm, with *k* = 3. This leaves us with three groups representing distinct coherent behaviors, labelled G1, G2, and G3, as well as the incoherent group, which we label *N*. These groups define the coloration of the top panel of Figure 5. Comparing these group memberships to those given by calculating the group polarization and rotation, *O*_*p*_ and *O*_*r*_, and classifying behavior as in [24] we find that group G1 contains 85.86% of the milling frames, G2 and G3 together comprise 87.47% of the polarized frames, while N holds 65.85% of the swarming frames (another 32.01% are relegated to G2). The full results of the classification appear in Table 1.

**Table 1.**
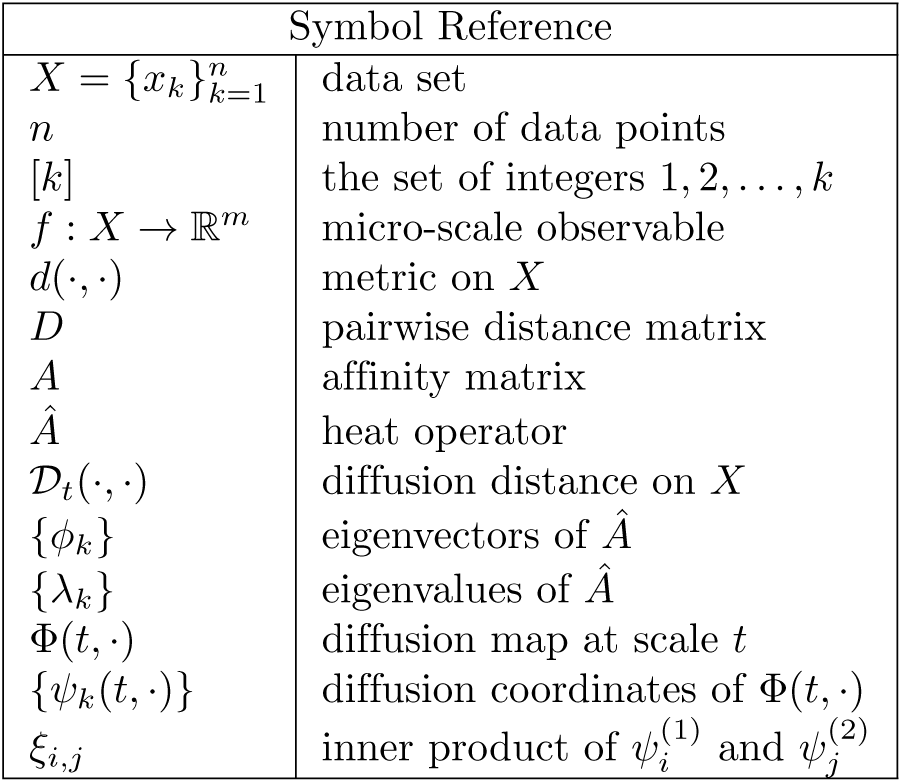
List of symbols used.

| Symbol Reference |  |
| --- | --- |
| $X = \{x_k\}_{k=1}^n$ | data set |
| $n$ | number of data points |
| $[k]$ | the set of integers $1, 2, \dots, k$ |
| $f : X \rightarrow \mathbb{R}^m$ | micro-scale observable |
| $d(\cdot, \cdot)$ | metric on $X$ |
| $D$ | pairwise distance matrix |
| $A$ | affinity matrix |
| $\hat{A}$ | heat operator |
| $\mathcal{D}_t(\cdot, \cdot)$ | diffusion distance on $X$ |
| $\{\phi_k\}$ | eigenvectors of $\hat{A}$ |
| $\{\lambda_k\}$ | eigenvalues of $\hat{A}$ |
| $\Phi(t, \cdot)$ | diffusion map at scale $t$ |
| $\{\psi_k(t, \cdot)\}$ | diffusion coordinates of $\Phi(t, \cdot)$ |
| $\xi_{i,j}$ | inner product of $\psi_i^{(1)}$ and $\psi_j^{(2)}$ |

**Table 2.**
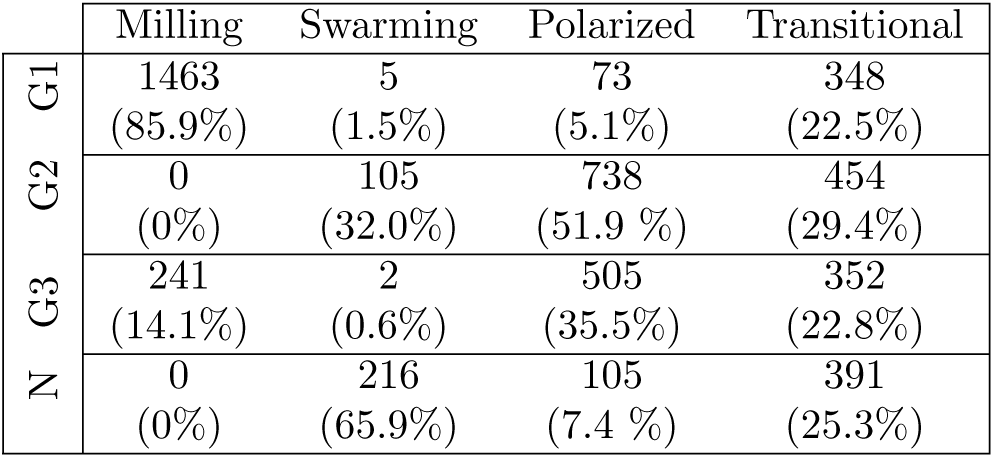
Counts of classifications by our map alignment-based statistics (rows) and by the macro-scale variable classification (columns) for the 5000 frames of fish movement data. The percentage of the frames of a given state (milling, polarized, etc.) which is found in each of our unsupervised groups (G1, G2, G3, N) are displayed below the counts.

|  | Milling | Swarming | Polarized | Transitional |
| --- | --- | --- | --- | --- |
| G1 | 1463<br>(85.9%) | 5<br>(1.5%) | 73<br>(5.1%) | 348<br>(22.5%) |
| G2 | 0<br>(0%) | 105<br>(32.0%) | 738<br>(51.9 %) | 454<br>(29.4%) |
| G3 | 241<br>(14.1%) | 2<br>(0.6%) | 505<br>(35.5%) | 352<br>(22.8%) |
| N | 0<br>(0%) | 216<br>(65.9%) | 105<br>(7.4 %) | 391<br>(25.3%) |

That the polarized frames are divided into two separate clusters suggests that there are two distinct modes of polarized behavior exhibited by the fish. Upon inspecting *ξ*(*t*) for the frames from each cluster, we find that the two groups are essentially distinguished by their relative levels of organization; for example, the mean value of *Z*_15_ for cluster G2 was 6.3186, while for G3 the mean map alignment statistic is 12.0878. Practically, the frames of cluster G2 typically consist of polarized group motion with multiple subgroups of fish following distinct paths; we call this *β*-polarized behavior. This creates a macro-scale division in the network structure of *Â*^(*v*)^ that is not present in the network structure of *Â*^(*x*)^, which leads to a lower alignment between their top eigenvectors; here (*ξ*_1,1_(*t*)) = 0.176. On the other hand, the frames of G1 feature a single, unified group motion, leading to a high alignment ⟨(*ξ*_1,1_(*t*)⟩ = 0.331) between the top diffusion coordinates of each network. We refer to this as *α*-polarized schooling behavior. It is possible that there is a correspondence between the empirically defined *β*-polarized activity and the *dynamic parallel* group behavior described in the family of models of [26].

One may approach the problem of classifying the covariance vectors {*ξ*(*t*)} in numerous ways; we chose the *k*-means algorithm for its simplicity. However, it requires the user to fix the number of desired clusters beforehand. As the prior work of [24] only defined two separate coherent behaviors, milling and polarized, one may wonder what results we would find if we instead restricted the classification to two clusters. We did this and, labelling the groups G1^*^ and G2^*^, found that G1^*^ successfully captures all the milling frames, and G2^*^ is predominantly composed of polarized frames (see Table S1). However, roughly 1 in 4 polarized frames are relegated to G1^*^. Table S2 shows how the two *k*-means clusterings apportion the frames. In particular, the *α*-polarized frames tend to be classified as milling. By increasing the number of subgroups to *k* = 3, we allow the algorithm to perform a more sensitive clustering, which improved the correlation between our groupings and those of [24], as well as distinguished between two forms of polarized behavior that were grouped together in that work.

## Discussion

To identify different states of collective behavior in complex systems, we have developed and applied a new methodology based on measuring changes in the mutual geometry of a given system as it is viewed through different lenses, using diffusion maps. The method is general and objectively produces system-specific, macro-scale variables, *Z*_*k*_ and *ξ* for tracking the onset of and distinguishing between different regimes of collective behavior. To test the method, we applied it to synthetic data produced from a well-known model of birds flocking, and to empirical data on fish schooling. In both cases the macro-scale quantities constructed using diffusion maps provided an accurate description of the system in terms of the degree and type of collective behavior present.

The analysis, while data-driven, does suggest methods of controlling complex systems, or nudging them towards a desired state, as in [27]. For example, the top panel of Figure 5 suggests that swarming behavior is typically bookended by a *β*-polarized state. Then to bring the school into a swarming state from an *α*-polarized state, one might attempt to force the fish to break into multiple polarized subgroups (by instituting an obstruction, say). Achieving a misalignment of the leading diffusion coordinates from the spatial- and velocity-based networks of the school will clearly impede any group consensus, making the school more likely to lose its coherence. This strategy is surely obvious in the case of schooling fish, but an analogous tactic could be employed to inform control over state change in markets or social networks as well.

Furthermore, the diffusion coordinates provide a quantitative basis for choosing how to align/misalign a system; one can use *in silico* experiments to identify changes to the network structure which would move the system towards the desired macro-state. Given a target covariance vector, the technique could perhaps be automated. One approach would be to perform spectral clustering of the network according to the present and target eigenfunctions. Agents belonging to the same cluster in both cases could be expected to remain well-connected during the transformation, simplifying the search for an appropriate perturbation to those that only increase/decrease the connectivity of those clusters. Reducing the problem space in this way would be a first step towards an online solution, advising system control in real time.

Flocking models and empirical data on fish schooling were studied here because the collective behavior from each system is well known. That is, the scales at which collective behavior emerges and the macro-scale variables that do a good job of describing group behavior are known. This allowed for a clear demonstration of the new methodology, though one can apply this framework to a variety of dynamic or static data sets, from subfields outside animal group dynamics, and areas other than biology. There are many non-biological systems, such as financial and housing markets, city-transit systems, and power-grid systems, which are known to exhibit emergent patterns at a range of scales, but for which the relevant micro-scale variables are less well known. In these cases one may compute map alignment in an uninformed fashion, testing a variety of metrics and variables, to discover latent emergent relationships.

In practice, the decision of which micro-scale metric to use may not always be clear, but the map alignment method offers a resilience to these particulars. In the two cases studied, familiarity with animal collectives and their emergent properties made the choice to use position and velocity data snippets and their respective metrics a straightforward decision. However, there are many other biological systems where the organizational variables to choose may not be so obvious [28]. While the exact variables controlling information propagation in such systems may be unknown, as demonstrated in [29], the underlying structure may still be recovered through measurements of different, related variables. That is, we take advantage of the gauge-invariance of the diffusion map approach. As many previous publications have pointed out [24, 30, 31], fish do not interact with one another based purely on their euclidean distance; other factors including rank of nearness (closest neighbors), relative alignment, and density. Therefore the metric *d*^(1)^ used above is a gross simplification of the role spatial organization plays. However, it captures enough of the true underlying organizational structure to allow us to detect emergent behavior.

It is worth noting, however, that the approach that we have developed has its own challenges relating to the nature of the data used. Diffusion maps inherently require a large number of data points for the diffusion operator to exhibit the regularity of the limiting operator on the manifold from which the data is, in theory, sampled. With the fish data though, the intermittent nature of the observations meant that the system size *n* varied between roughly 70 and 280 agents, and still the method proved resilient to these fluctuations.

Also, if one employs the GkNN kernel as we do here (see (1)) the scale-free property simplifies the task of resolving the macro-scale structure of the data; in particular, one does not need to adjust the kernel if the distances involved globally grow or shrink. But this entails a loss of information. As an example, if the system becomes highly organized when observed via the metric *d* (e.g. *d* corresponds to the velocity-based metric *d*^(2)^ above, and the fish all move as one) the corresponding data cloud (*X, d*) collapses towards a point mass. However, the GkNN kernel rescales distances in order to resolve their structure across scales, i.e. the rescaled distances only have meaning by virtue of their relative size. As a result the network’s features can become a measurement of the lower-order stochastic fluctuations of individual agents’ trajectories, rather than meaningful organization. This conflation of extremely organized and unorganized states could be overcome by tracking the mean inter-agent distance, for example, and may be important in application.

Measuring the map alignment between different networks induced by separate metrics on a graph serves as an entry point for several other possible analyses. One could use a set of training data to perform a classification of the various emergent behaviors of the system, then measure the stability of the system (risk of state change) by the current distance from *ξ*(*t*) to representative subgroups from the various states. For example, if the distance from *ξ*(*t*) to Group 1 is growing, and the distance to Group 2 is shrinking, one might anticipate a shift in macrostate. In this way one may estimate the risk of emergence, dissolution, or a large-scale shift of collective behavior. The authors believe, based on preliminary analysis of the fish schooling data, that such an approach could provide a new early warning metric for complex systems whose relevant macrovariables may be unknown, inhibiting the application of other early warning signals such as critical slowing down.

## Conclusion

The method that we have developed is a new data-driven approach for detecting cross-scale emergent behavior in complex systems. It quantifies the dependence present between various micro-scale variables (those exhibited at the agent-level), and formulates a signature of the macro-scale behavior exhibited by the collective. The method can also detect the onset or loss of organization in an unsupervised fashion. With sensible choices of micro-scale variables, which are now measured routinely as Big Data, any complex system can be studied in this way. Doing so presents new opportunities for studying complex systems through analyzing their changing geometry.

## Supporting information

### Idealized Simulations of Bird Flocking

Let *N* agents be initiated at time 0 with position chosen uniformly at random on the torus 𝕋= *S*^1^ × *S*^1^, and with bearing chosen uniformly at random from *S*^1^. Write *B*_*i*_(*r*) = {*j* : ‖*p*_*i*_ − *p*_*j*_‖ < *r*} for the set of agents within a radius *r* of agent *b*_*i*_. The dynamics of our flocking model are written as follows:

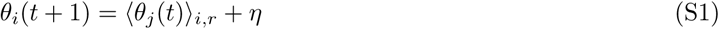

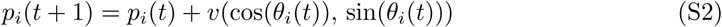

where ⟨*θ*_*j*_(*t*) ⟩_*i,r*_ is the mean bearing of the agents within *B*_*i*_(*r*) and *η* is a Gaussian noise with standard deviation *ϵ* > 0. (Note that this mean bearing is calculated by finding the local mean velocity, then setting ⟨*θ*_*j*_(*t*) ⟩_*i,r*_ equal to its angle with the abscissa.) Thus each agent’s position is updated by flying a distance *v* in the direction determined (up to the perturbation *η*) by the average of the directions of all other agents within *r* units of the focal agent. Allowing *r* to vary over time, transitions between flocking and disordered behavior can be observed. Figure 1 in the main text shows typical system states with *N* = 800 agents in the incoherent and coherent phases. Code for this simulation can be found below.

In the figures and analysis of the manuscript, the datasets are generated with *N* = 500, taking the noise to be *ϵ* = *π/*5, velocity *v* = 1*/*320, with minimum and maximum values of the interaction radius given by *r*_0_ = 0.0065 and *r*_1_ = 0.06.

### Empirical Data on Collective Behavior in Fish

The fish behavioral data was gathered by Tonstrøm et al. by recording video at 30 frames per second of a group of 300 golden shiners moving about a tank of shallow water. The water level was low enough to essentially limit the fish to planar motion. Image analysis was employed to locate the fish within each frame, then match them across frames when possible. This allowed the research team to associate a unique identifier with each fish while its trajectory was under observation.

These fish are highly social, and due to the difficulty of rendering individual fish when they are packed tightly into a region, and the possible conflation of two separate fish swimming near one another, individual fish are often ‘dropped’ from the data set. When this occurs the identifier associated with that fish is retired, and the position and velocity data for that particular trajectory terminates. However, when the fish again exceeds the liminal level of the image analysis it is registered as a ‘new’ fish and given a new identifier. As a result, the data garnered from a typical frame records position and velocity information for only 60 - 80% of the members, and individuals are typically tracked for about 200 frames (6.7 seconds) before its image is ‘lost’ and its identifier removed from future data. The resulting data set consists of 100298 frames (about 56 minutes) of position and velocity data for the school. For the study in the manuscript we selected a 5000 frame subset containing roughly an equal number of frames devoted to each of the behaviors observed (frames 40001 to 45000). This enabled us to use the simple *k*-means clustering algorithm, which otherwise suffers when the subgroups to be identified have widely varying sizes. (This is because the classical *k*-means algorithm creates hyperspherical clusters, where the hyperspheres must be centered within the convex hull of the data and have equal radii. Smaller clusters necessitate a description via smaller hyperspheres. See [32].)

As discussed in [25], human observation of the schooling behavior identified three separate behavioral regimes for the fish: milling, swarming, and polarized (see Figure S1). In the milling state, the fish rotate as a group, circulating about a fixed point; this is a highly coherent group state. While swarming, the fish trajectories are disordered, with very little discernible group alignment. In this state the school is stationary, though individual fish may be moving in an uncoordinated fashion. Last, the second coherent group structure is the polarized state, which is characterized by the group having a coherent direction of motion, and strong local alignment among the agents. As with the flocking model above, the variables of interest are the two-component vectors of fish position, *p*_*i*_, and fish velocity, *v*_*i*_.

### Metrics

Let *x*_*i*_ ∈ *X* be the set of observations for an agent (either a particle in the flocking simulation or a golden shiner found in the school) at a fixed time *t*, while *p*_*i*_(*t*), *θ*_*i*_(*t*), and *v*_*i*_(*t*) are the *i*^th^ agent’s position, heading, and velocity, respectively. The first metric, *d*^(1)^, is simply the euclidean distance between the agents,

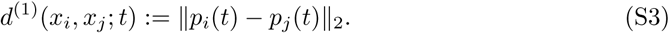

For the flocking simulation, the second metric is the following

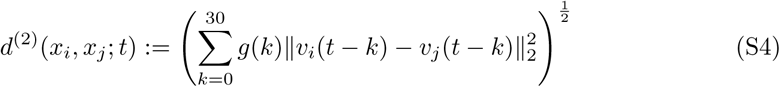

where *v*_*i*_(*t*) = *v*(cos(*θ*_*i*_(*t*)), sin(*θ*_*i*_(*t*)) is the *i*^th^ velocity at time *t*. The smoothing term *g*(*k*) is proportional to exp (−*k*^2^*/*30), but normalized so that ∑_*k*_ *g*(*k*) = 1. That is, *d*^(2)^ is a gaussian smoothed time-average of the *L*^2^ norm between velocities over the past 30 time steps.

For the fish, we consider a generalization of the previous expression for *d*^(2)^. Rather than comparing the two fishes’ heading time series at simultaneous moments, we allow one time series to be shifted into the past by a lag of up to *L* = 60 video frames, or two seconds.

Define

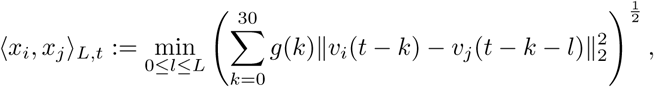

where *g*(*k*) is again a gaussian smoothing kernel as in the previous paragraph. Thus *l* represents a lag allowing us to compare not only concurrent velocity profiles over 30 steps, but also cases where fish *i* is mimicking a trajectory of fish *j* up to two seconds in the past. Note that ⟨*x*_*i*_, *x*_*j*_⟩_*L,t*_ ≠ ⟨*x*_*j*_, *x*_*i*_⟩_*L,t*_, as the righthand term calculates the distance between the trajectories under the assumption that *j* is following *i*. We define our distance as

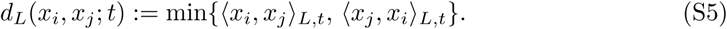

**Table S1.** Counts of classifications by our map alignment-based statistics (rows) and by the macro-scale variable classification (columns) for 5000 frames of fish movement data. The percentage of the frames of a given state (milling, polarized, etc.) which is found in each of our unsupervised groups (G1^*^, G2^*^, N) are displayed below the counts.

|  | Milling | Swarming | Polarized | Transitional |
| --- | --- | --- | --- | --- |
| G1* | 1704<br>(100.0%) | 6<br>(1.8%) | 375<br>(26.4%) | 606<br>(39.2%) |
| G2* | 0<br>(0%) | 106<br>(32.3%) | 941<br>(66.2 %) | 548<br>(35.5 %) |
| N | 0<br>(0%) | 216<br>(65.9%) | 105<br>(7.4 %) | 391<br>(25.3%) |

**Table S2.**
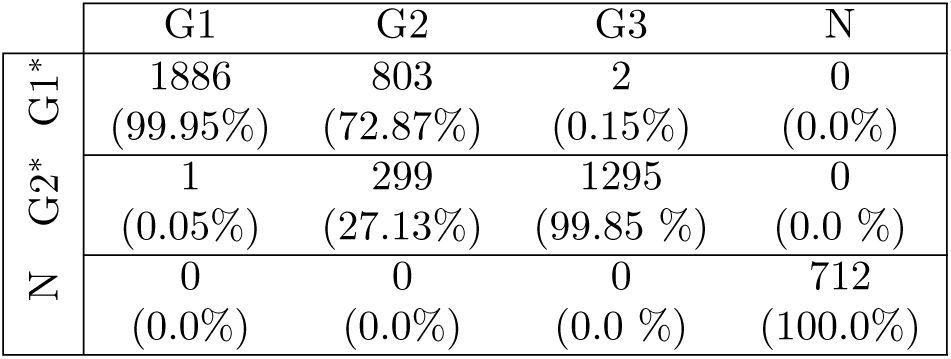
Comparison of the two *k*-means classifications for 5000 frames of fish movement data. The 2-group clustering gave G1^*^ and G2^*^, corresponding to frames of high and low macro-scale organization, respectively. The 3-group clustering gave G1, G2, and G3, corresponding to milling, *α*-polarized, and *β*-polarized behaviors, respectively. The percentages are with respect to all the frames in the entry’s column.

**Fig S1.**
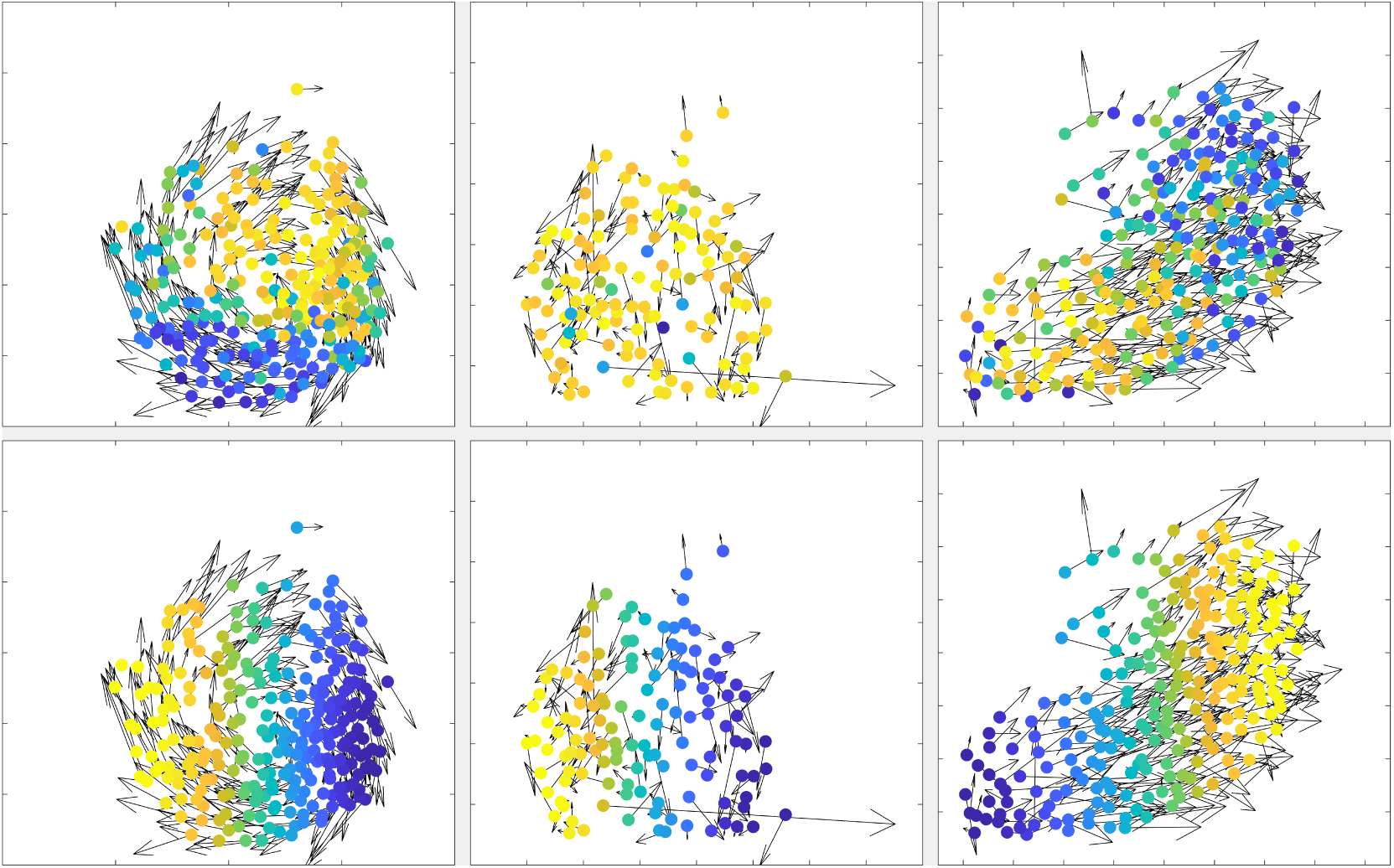
Plots of fish position (colored markers) and velocity (arrows) at three example times, colored by diffusion map coordinates 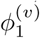 [top row; i.e. produced from analyzing correlations in velocities] and 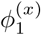 [bottom row; i.e. produced from analyzing spatial proximity]. The emergent behaviors from left to right are milling, swarming, and polarized motion.

## Supplementary Movies

**Movie 1:** A sample run of the flocking model (*ϵ* = *π/*5, *v* = 1*/*320) where the interaction radius *r*(*t*) is equal to *r*_0_ = 0.0065 until time *t* = 800, is equal to *r*_1_ = 0.06 for times after *t* = 1200, and increases linearly from *r*_0_ to *r*_1_ over time steps 800 to 1200. Each subplot is colored according to one of the top two diffusion coordinates, as labeled. The top two subplots display the first and second coordinates from the *d*^(1)^-derived affinity matrix (see equation (S3)), while the bottom subplots are colored by the diffusion coordinates from the *d*^(2)^-based affinity matrix (see equation (S4)).

**Movie 2:** A sample run of the flocking model (*ϵ* = *π/*5, *v* = 1*/*320) where the interaction radius *r*(*t*) varies sinusoidally between *r*_0_ = 0.0065 and *r*_1_ = 0.06, with the period of oscillation increasing linearly for the duration of the time series: *r*(*t*) = 0.0. Each subplot is colored according to one of the top two diffusion coordinates, as labeled. The top two subplots display the first and second coordinates from the *d*^(1)^-derived affinity matrix (see equation (S3)), while the bottom subplots are colored by the diffusion coordinates from the *d*^(2)^-based affinity matrix (see equation (S4)).

**Movie 3:** This video displays a subset of the fish schooling data used, colored by various diffusion coordinates. We restrict the frames displayed to keep the file size reasonable while still showcasing the various modes of behavior described in the main text. The time step displayed in each frame corresponds with the time steps shown on the abscissa of Figure 5. The top two subplots display the first and second coordinates from the *d*^(1)^-derived affinity matrix (see equation (S3)), while the bottom subplots are colored by the diffusion coordinates from the *d*^(2)^-based affinity matrix (see equation (S5)).

## Acknowledgments

The authors would like to acknowledge support from the DARPA YFA project N66001-17-1-4038 and Kolbjørn Tunstrøm for providing the fish schooling data

